# Next generation of anti-PD-L1 Atezolizumab with better anti-tumor efficacy *in vivo*

**DOI:** 10.1101/2020.06.30.166207

**Authors:** Maohua Li, Rongqing Zhao, Jianxin Chen, Wenzhi Tian, Chenxi Xia, Xudong Liu, Yingzi Li, Yuyuan Yan, Song Li, Hunter Sun, Tong Shen, Wenlin Ren, Le Sun

## Abstract

Some cancer patients treated with Atezolizumab, PD-L1 antibody drug launched by Genentech, quickly developed anti-drug antibody (ADA), led to loss of efficacy. This was likely due to the heavy aggregation of Atezolizumab, caused by mutation of N297A for removing unwanted antibody-dependent cytotoxicity (ADCC) of IgG1 antibody drug. Here, we developed a new version of Atezolizumab (Maxatezo), which was demonstrated better anti-tumor efficacy *in vivo*. In Atezolizumab, we mutated 297A to 297N back to bring back the glycosylation, and inserted a short sequence GGGS between G237 and G238 in the hinge region of the IgG1 heavy chain. Our data shown that insertion of GGGS, without altering the anti-PD-L1 antibody affinity and inhibitory activity, completely abolished the ADCC activity, as same as Atezolizumab. Moreover, the insertion of GGGS, without altering the glycosylation profile of IgG1, increased the yields of anti-PD-L1 antibody considerately. Additionally, glycosylation improved the stability yet reduced the amounts of aggregations in the antibody solutions. In turn, the level of ADA in animals treated with Maxatezo was 70% lower than the ones treated with Atezolizumab. Most importantly, at the same 10mg/kg dose, the anti-tumor activity of Maxatezo had attained 98% compared to that of Atezolizumab at 68%.

## Introduction

In recent years, with the deepening of the research on the mechanism of tumor immune escape and tumor microenvironment, immunocheckpoint inhibitor based therapies have shown satisfactory clinical efficacy in a variety of tumors. Immunotherapy play an important role in the treatment of malignant tumors. And the use of anti-PD-1 and its ligand PD-Ll antibodies in the treatment of malignant tumors has become a focus of research. Several anti-PD-1 or PD-L1 antibodies were approved by FDA, such as Pembrolizumab of Merck, Nivolumab of BMS and Atezolizumab of Roche. In 2019, the sale of therapeutic antibodies, which target PD-1/PD-L1, is 22.9 billion U.S. dollar. However, regardless of the sales of Pembrolizumab ($11.084 billion) and Nivolumab ($8.017 billion), the sales of Atezolizumab is only $2.091 billion. The deglycosylation of the antibody resulting in the reduced stability, formation of aggregation, which can significantly increased immunogenicity, and more intense production of anti-drug antibody (ADA). In the expansion of indications, due to its immunogenicity, a number of clinical Phase III did not reach the clinical end point (in 2017, IMvigor211 for advanced bladder cancer failed in clinical Phase III; in 2018, IMblaze370 for Colorectal Cancer failed in clinical Phase III; in 2019, IMspire170 for Melanoma failed in clincal Phase III)^1^, and the approved indications were far less than those of rival products in the same field. Therefore, in terms of market occupancy and annual sales, it lags far behind Keytruda and Opdivo.

It is well-known in antibody manufacturing that incomplete glycosylation will lead to aggregations of antibodies^2^, which in turn will induce strong ADA in treated patients^3^. Aglycosylation of antibody made the cases even worse. Atezolizumab’s drug label includes a warning to not shake the vial or discard the drug if the solution turns cloudy. In its pre-clinical and clinical studies, drug treated monkeys developed ADA at a 100% rate, while in cancer patients whose immune systems had been severely damaged by chemo or radiotherapy, ADA was developed at a 41.5% rate^4^. The fast development of neutralizing ADA forced the dosage of Atezolizumab to be escalated to a record high of 1200 mg per injection^4^ and still failed to reach end-points of several Phase III clinic trials^1^.

Therapeutic antibodies have different mechanisms of action, such as 1) neutralizing antibodies that block the target/pathogen’s biological activities, 2) clearing antibodies mediated by antibody-dependent cell phagocytosis (ADCP) to remove the target/pathogen from the body, or 3) targeting antibodies via antibody-dependent cytotoxicity (ADCC) to recruit natural killer (NK) cells and other effector T cells to kill the pathogens or tumor cells. Based on the intended mechanisms of action, researchers and drug developers may choose to utilize different isotypes.^5, 6^. Most antibody drugs choose IgG1, IgG2 and IgG4 isotypes, while IgG3 is avoided due to its instability. The sequence homology between IgG1, IgG2 and IgG4 is more than 90% with the primary differences resting within the hinge region and CH2 domain, which contain the binding sites for different FcγRs^7, 8^.

FcγR engagement is essential for the Fc functions of IgGs ^9, 10, 11^. Binding to antigens by the antibody can change the conformation of the Fc region to expose the binding sites for FcγRs, which in turn can activate ADCC and/or ADCP activity ^12, 13^. The human FcγR family consists of the activating receptors FcγRI, FcγRIIA, and FcγRIIIA, and the inhibitory receptor FcγRIIB^14^. FcγRI and FcγRIIA are expressed by macrophages, and involved in ADCP function^15, 16^, and FcγRIIIA which is expressed on NK cells is important for ADCC function^17^.

For some antibodies targeting specific antigens on the surface of tumor cells, ADCC mediated effects allow NK cells to effectively kill tumor cells^18^. For development of these types of therapeutic antibodies, IgG1 isotypes with strong ADCC functions are preferred, and some of the antibodies are even modified to further enhance their ADCC effects^19^.

For blocking antibodies, such as antibodies targeting soluble cytokines (TNF alpha and IL17A) or some immune checkpoints (CTLA-4, PD-1, PD-L1), the ability to bind to FcγR is not desirable and the cytotoxicity brought about by ADCC/ADCP should be prevented. The binding affinity of IgG1 to FcγRs on effector cell surfaces is highly dependent on the N-linked glycan at asparagine 297 (N297) in its CH2 domain^20, 21^, with a loss of binding to the FcγRs observed in N297A point mutants^22, 23^, enzymatic Fc deglycosylation^24^, recombinant IgG expression in the presence of the N-linked glycosylation-inhibitor tunicamycin^25^, or expression in bacteria^26, 27^. In addition, the nature of the carbohydrate attached to N297 modulates the affinity of the FcγR interaction as well^28, 29^. Aglycosylation of IgG1 has also been used to completely remove the unwanted ADCC/CDC^23, 30^. FDA-approved anti-PD-L1 antibody drugs Atezolimumab is human IgG1 without glycosylation by N297A mutation.

Structures of IgG1 Fc region show that the oligosaccharide attached to N297 is hidden in the cavity between the CH2 domains from the two heavy chains. When antibodies bind to the antigen, there are conformation changes in the Fv regions, which result in a domino effect via the hinge region to the remaining Fc region, leading to a cascade of conformation changes including the exposure of oligosaccharides attached to the N297 and the formation of the FcγR binding domain. Then, the antigen-attached antibody would bind to the FcγR on the surfaces of effector cells, leading to ADCC activation^21^. In theory, if one could block the conformation change signal transduction from the Fv region to the Fc region, this should also prevent the formation of the FcγR binding site, and thus prevent the ADCC activation.

In this paper, based on the knowledge of an antibody’s 3D conformation changes post antigen binding, we propose a novel way to design antibody drugs without ADCC function by simply inserting a flexible amino acid sequence such as GGGS into the hinge region of the antibody’s Fc region. Our data shown that such an insertion completely abolished the antibody’s ADCC. The effects on binding affinity (EC50), inhibitory activity (IC50), the glycosylation profile, expression level, stability, immunogenicity and anti-tumor activity were also examined.

## Results

### Design and gene synthesis of IgG1 Fc for removal of Fc functions of anti-PD-L1 antibody

Based on the structural information acquired from different antibodies in Protein data bank (e.g., PDB ID: 1IGT), we hypothesized that an insertion of a short but very flexible sequence in the hinge region or somewhere upstream of the glycosylation site of N297 may cut off the stress transmission signal between the Fv and Fc domain. In this study, short sequence of GGGS was chosen and inserted between G237 and G238 of human IgG1 heavy chain. The sequence of GGGS has been used in many approved biological drugs, such as scFV and Fc-fusion proteins, as flexible linkers without any known adversary effects or new immunogenicity in patients. For reverse engineering of Atezolizumab, a back mutation of A297N was also introduced into the original heavy chain to restore the glycosylation (Fig. S1).

Genes encoding either the heavy chains of Atezolizumab or the modified version Maxatezo were chemically synthesized and ligated into expression vectors. Transient co-transfections of the light chain and the heavy chain of IgG1 were carried out in CHO-K1 cells. The recombinant antibodies were purified out from the culture supernatants and subjected to further characterizations.

To evaluate whether there will be any negative impact on antibody affinity, we compared their affinities to the recombinant PD-L1 using an indirect ELISA. As shown in Fig. 1A, both Atezolizumab and Maxatezo bind to PD-L1 in a dose-dependent manner, with EC50 between 0.22∼0.32 nM. No negative impact on antibody affinity by insertion was observed.

**Figure 1.**
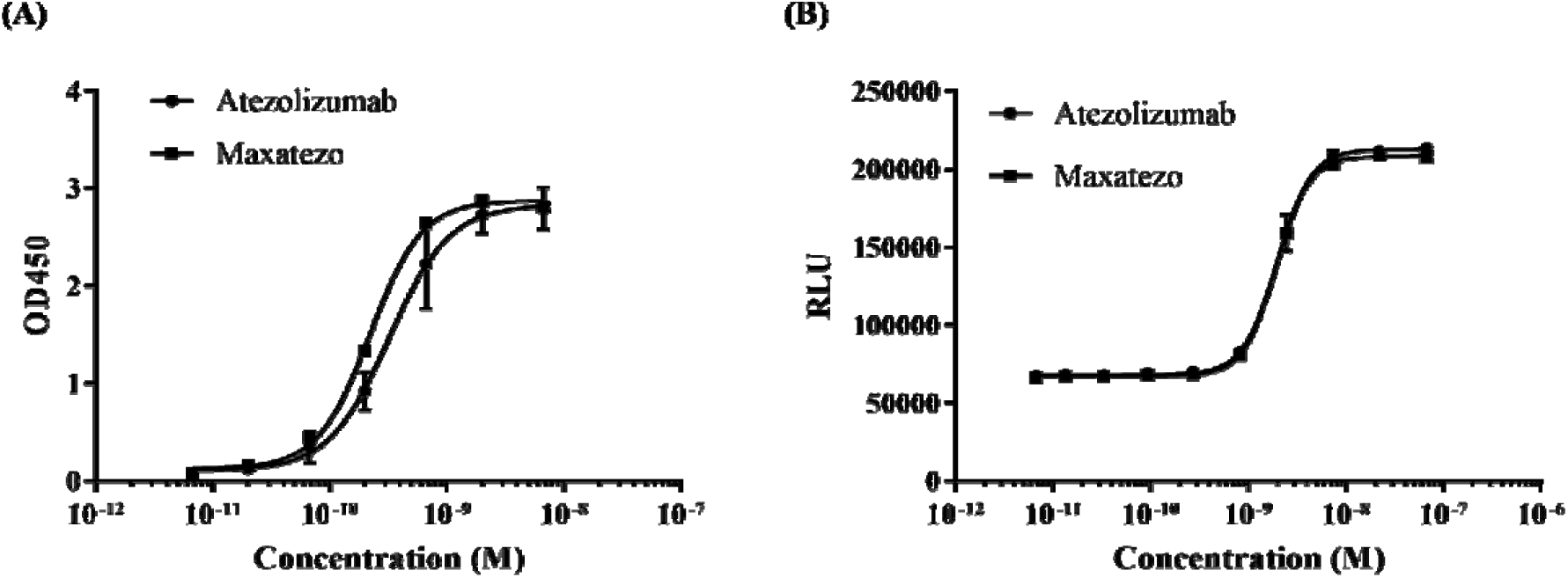
The Fc function removed PD-L1 antibody, Maxatezo. (a). Binding affinities of Maxatezo and Atezolizumab, measured by Indirect ELISA against recombinant PD-L1. The square mark represents Maxatezo, and the circular mark represents Atezolizumab. The horizontal coordinate is antibody concentration and the vertical coordinate is absorbance value at 450nm. (b). Inhibitory affinity evaluation of Maxatezo and Atezolizumab, measured by *in vitro* cell-based Bioassay using Jurkat-PD-1-NFAT cell line that stably expresses PD-1 with the reporter luciferase gene. The square mark represents Maxatezo, and the circular mark represents Atezolizumab. The horizontal coordinate is antibody concentration and the vertical coordinate is mean value of the relative fluorescence units (RLU).

Next, we examined the potential impact of the insertion on inhibitory activities of anti-PD-L1 antibodies using an *in vitro* co-culture of Jurkat-PD-1-NFAT cell line that stably expresses PD-1 with the reporter luciferase gene and CHO-PD-L1-CD3L cells that stably expresses PD-L1 and a membrane-anchored anti-CD3 single chain antibody fragment (scFv). The binding of PD-1 on the surface of Jurkat-PD-1-NFAT cells to the PD-L1 on the CHO-PD-L1-CD3L cells can block the downstream signal transduction of CD3 activated by the binding of anti-CD3 scFv, thus inhibiting the expression of luciferase in Jurkat-PD-1-NFAT cells. When PD-1 antibody or PD-L1 antibody is added, it will reverse the inhibition of luciferase expression. Addition of either Maxatezo or Atezolizumab reversed the inhibition of PD-1 on CD3-activated luciferase expressions in a dose-dependent manner, and both had IC50 between 1.95∼1.99 nM (Fig. 1B).

Clearly, the insertion of GGGS between G237 and G238 of human IgG1 heavy chain showed no significant negative impact on antibody’s affinity and inhibitory activity.

Next we studied whether the insertion indeed removed the ADCC of human IgG1 or not. Three anti-PD-L1 antibodies, the IMM25 (ADCC-enhanced version by afucosylation of N297), Atezolizumab (De-ADCC version by N297A mutation) and Maxatezo (De-ADCC version by GGGS insertion) were tested in parallel for their ADCC activities using a co-culture cell killing assay *in vitro*. Briefly, CFSE-pre-dyed PD-L1 overexpressing Raji cells were mixed with FcR overexpressing TANK cells in the presence or absence of PD-L1 antibodies. Once the PD-L1 antibody binds to the PD-L1 on the surface of the Raji cells, the FcR of the TANK cells will bind to the PD-L1-bound anti-PD-L1 antibody, which in turn will activate the releasing of cytotoxic cytokines from the TANK cells and eventually kill the Raji cells.

The positive control, an ADCC-enhanced anti-PD-L1 antibody IMM25 had very strong ADCC activity in a dose-dependent manner, while both Maxatezo and Atezolizumab showed no ADCC activities at all (Fig. 2). It indicates that GGGS insertion in the Fc hinge region completely abolished the ADCC activity of human IgG1.

**Figure 2.**
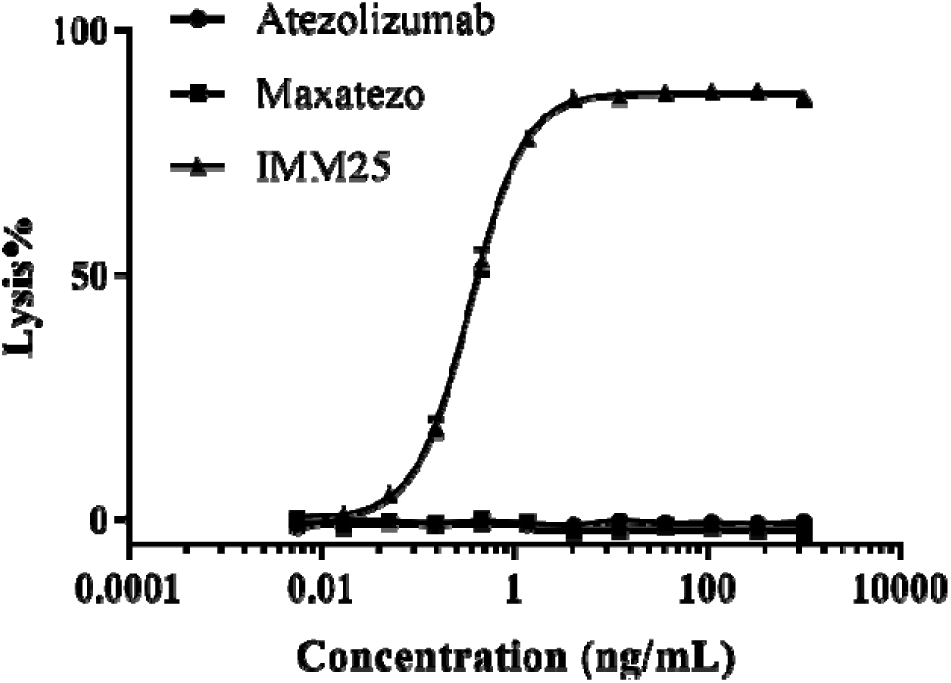
ADCC activities of three different anti-PD-L1 antibodies. The ADCC activities of the different PD-L1 antibodies were measured by an *in vitro* cell killing assay using PD-L1 overexpressing Raji cells together with FcR overexpressing TANK cells in the presence or absence of PD-L1 antibodies. The line with triangles represents the IMM25 (ADCC enhanced anti-PD-L1 antibody), the line with squares represents the Maxatezo, (anti-PD-L1 antibody with GGGS insertion), the line with solid circles represents the Atezolizumab (anti-PD-L1 antibody with N297A). The horizontal coordinate is antibody concentration and the vertical coordinate is the percentage of ADCC induced cell lysis.

### Effects on antibody’s expression, glycosylation, stability and immunogenicity

For therapeutic application, the recombinant antibody needs to be produced using a stable mammalian cell system at reasonable high expression level. Stable expression CHO cell pools were established for both Atezolizumab and Maxatezo. Two production runs for each antibody were carried out using 2 L and/or 5 L Bioreactors with same manufacturing parameters. On day 11, the culture supernatants were collected and the antibody concentrations in the culture supernatants were measured. As shown in Table 1, the expression levels of Maxatezo by stable CHO cell pools were between 4.07∼4.39 g/L, which qualified the industry expectation of 4 g/L or higher. However, the expression levels of Atezolizumab were approximately 1.36 g/L, much lower than that of Maxatezo. Possibly, aglycosylation of Atezolizumab could lead to in-correct folding of antibodies, which in turn cause the decrease of secretion. After cloning and selection, the expression level of Maxatezo by CHO monoclonal cell line increased to 6.2 g/L. Our data had suggested that insertion of GGGS does not have negative impact on recombinant antibody expression. Adding back of the glycosylation actually help to improve the production level.

**Table 1.** Expression of antibodies by stable CHO cell pools

| Run # | Culture Volume | Antibody Conc. (g / L) |
| --- | --- | --- |
| **B01 Atezolizumab** | 5L | 1.36 |
| **B02 Atezolizumab** | 2L | 1.30 |
| **B03 Maxatezo** | 2L | 4.39 |
| **B04 Maxatezo** | 2L | 4.07 |

Note: The stable cell pools expressing either Atezolizumab or Maxatezo were seeded in Bioreactors and the culture supernatants were harvested on Day 11. The antibodies in the culture supernatants were separated by analytic HPLC and the antibody concentrations were determined based on the areas of the peaks at OD 280.

High molecular weight (HMW) aggregations of antibody drugs are the major causes of anti-drug antibody (ADA) ^3^. Aglycosylation of IgG1 could make the aggregation even worse. To examine the levels of aggregates in the final drugs, Maxatezo and Atezolizumab were separated by HPLC-SEC. The percentages of HMW in the final drugs were estimated to be less than 1.0%, based on the peak areas of monomer and HMW, seems met the quality requirement for antibody drug (Fig.3, Table 2).

**Table 2.**
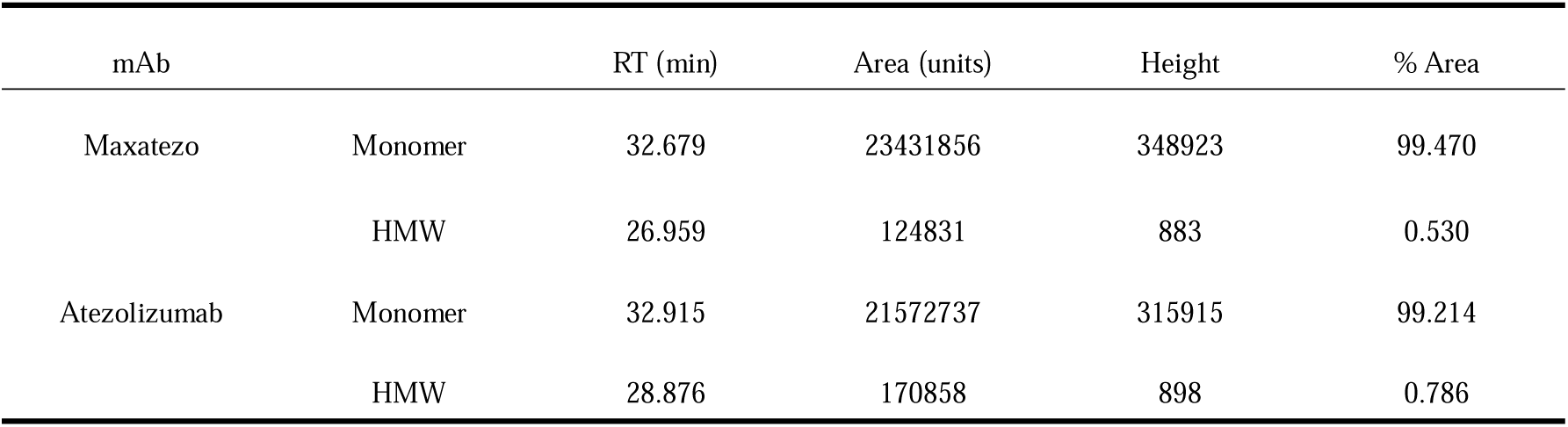
HPLC-SEC Analysis

| mAb |  | RT (min) | Area (units) | Height | % Area |
| --- | --- | --- | --- | --- | --- |
| Maxatezo | Monomer | 32.679 | 23431856 | 348923 | 99.470 |
|  | HMW | 26.959 | 124831 | 883 | 0.530 |
| Atezolizumab | Monomer | 32.915 | 21572737 | 315915 | 99.214 |
|  | HMW | 28.876 | 170858 | 898 | 0.786 |

**Fig 3.**
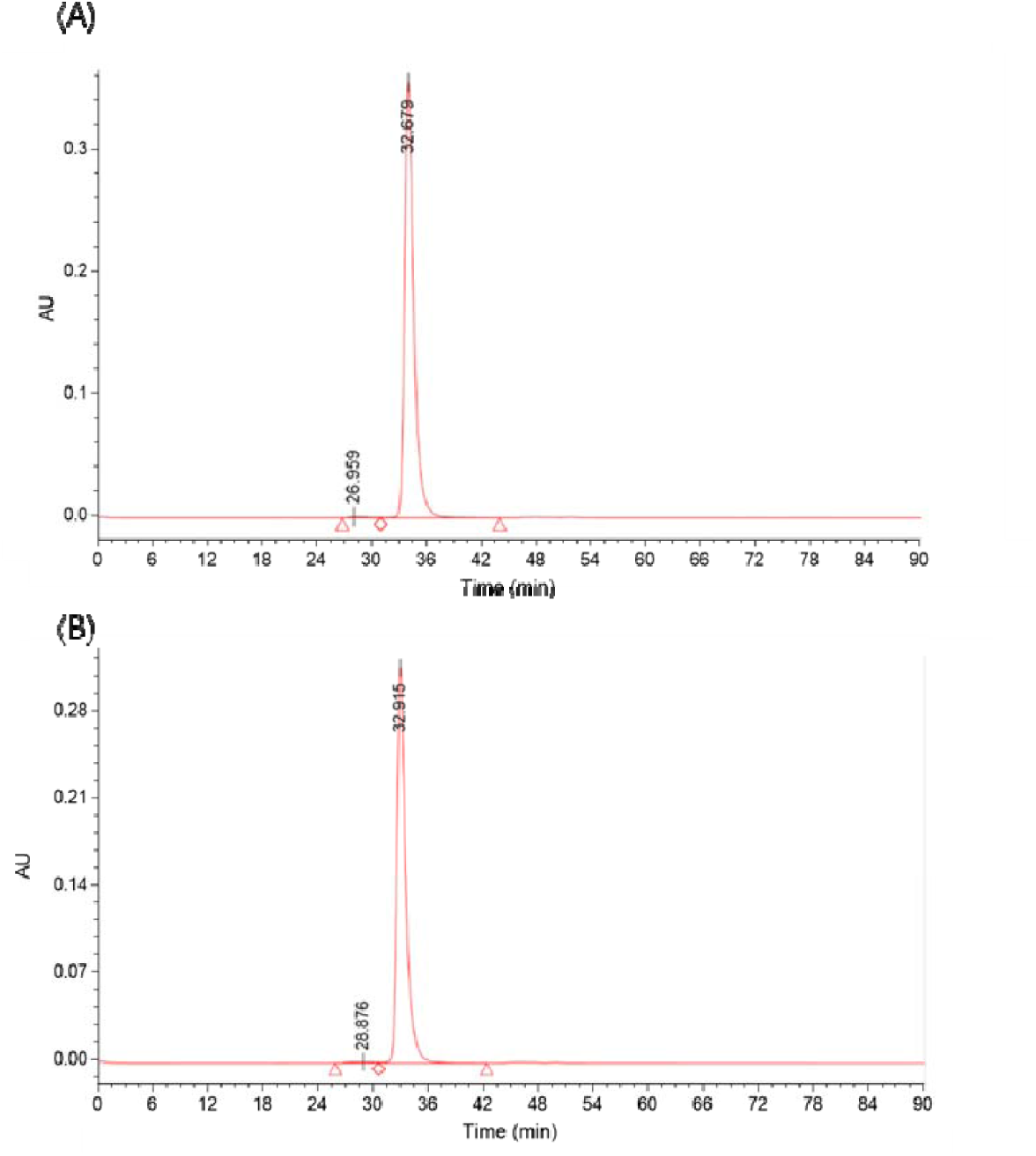
Purity analysis of Maxatezo & Atezolizumab by HPLC-SEC. (A) The gel filtration chromatography of Maxatezo; (B) The gel filtration chromatography of Atezolizumab. The horizontal coordinate is time and the vertical coordinate is absorbance value at 280nm.

However, when we took a close look at the areas for the Monomers, Atezolizumab had absorbance of 21,572,737 units, 10% less of protein than Maxatezo’s 23,431,856 units, even though the loading amounts were the same. It is possible that Atezolizumab has significant amount of large aggregation which were removed by the pre-filter of the HPLC columns.

Not only glycosylation is important for the correct conformation, secretion and stability of protein, its carbohydrate moiety also plays great role in its affinity to its receptor(s). Therefore, it is important to confirm the insertion does not result significant changes to the profile of the glycosylation. Both Maxatezo and Atezoliumab were subjected to digestion with PNGase F to release the glycan from the antibodies. The enzymatic products were labeled with 2-Aminobenzamide (2-AB), then separated by Hydrophilic Interaction Chromatography (HILIC) with a fluorescence detector. While there was no glycan detected in the sample digested from Atezoliumab, the glycans with different sizes were observed in the sample prepared from Maxatezo (Fig. S2).

The glycan profiles of different clones of Maxatezo were further analyzed. As shown in Table 3 and Figure 4, all clones exhibited similar profiles of glycan compositions and the distribution of the glycoforms appeared to be normal. Our data indicates the insertion of GGGS does not change the glycan profile of human IgG1.

**Table 3.** Glycosylation profile of Maxatezo

| Clone ID | G0-GN (%) | G0F-GN (%) | G0 (%) | G0F (%) | Man-5 (%) | G1F (%) | G1F' (%) | G2F (%) | Others (%) |
| --- | --- | --- | --- | --- | --- | --- | --- | --- | --- |
| **C9** | 0.1 | 0.4 | 2.5 | 72.8 | 0.7 | 13.2 | 5.0 | 1.5 | 3.8 |
| **C10** | 0.2 | 0.5 | 2.6 | 70.6 | 0.8 | 14.0 | 5.3 | 1.6 | 4.4 |
| **C11** | 0.2 | 0.5 | 4.5 | 68.7 | 1.7 | 13.3 | 5.1 | 1.3 | 4.7 |
| **C12** | 0.1 | 0.5 | 2.4 | 73.8 | 0.7 | 12.6 | 4.8 | 1.3 | 3.8 |
| **C13** | 0.3 | 0.9 | 2.9 | 62.7 | 2.5 | 17.3 | 6.1 | 2.5 | 4.8 |
| **C14** | 0.1 | 0.5 | 2.6 | 71.9 | 0.7 | 13.5 | 5.1 | 1.5 | 4.1 |
| **C15** | 0.1 | 0.5 | 2.4 | 70.0 | 0.8 | 14.4 | 5.6 | 1.6 | 4.6 |
| **C16** | 0.2 | 0.5 | 2.6 | 71.7 | 0.7 | 13.5 | 5.1 | 1.5 | 4.2 |
| **C17** | 0.3 | 0.9 | 3.1 | 63.4 | 2.2 | 16.9 | 6.1 | 2.4 | 4.7 |
| **C18** | 0.2 | 0.5 | 2.6 | 70.0 | 0.7 | 14.6 | 5.6 | 1.7 | 4.1 |

**Figure 4.**
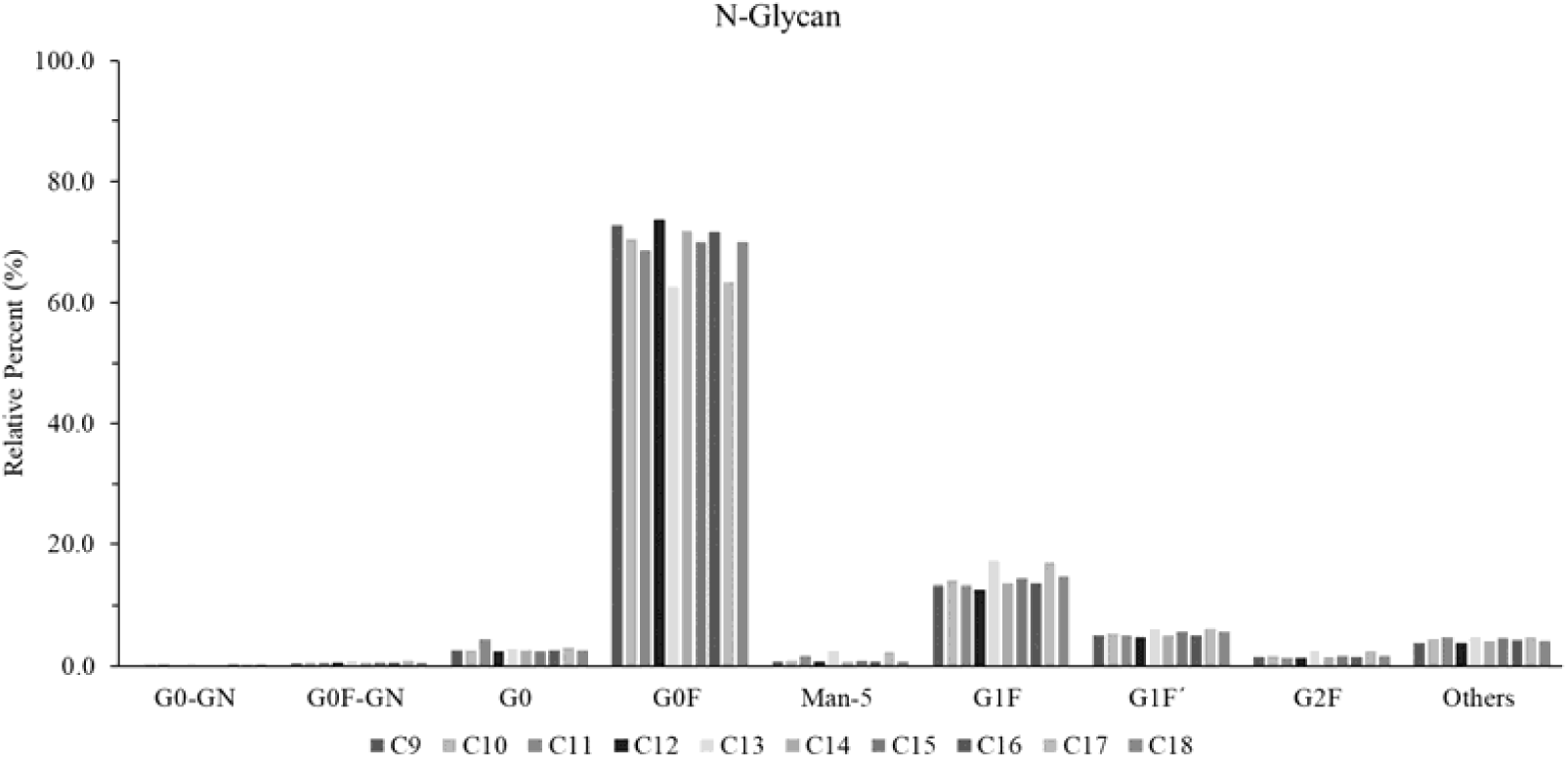
N-glycan isotype evaluation of Maxatezo. The relative percent of different glycan was verified. The horizontal coordinate is glycan isotype and the vertical coordinate is relative percent.

Thermal stability is one of the key factors for antibody drug development and it is highly influenced by the status of glycosylation^2^. To evaluate the stability of Maxatezo and Atezolizumab, both antibodies were heated at 60 □ for 10 minutes, then centrifuged at 10000 rpm for 5 minutes, and filtered using 0.2μm filters. Samplings were taken at each step and the protein concentrations were measured at OD_280_ using a UV spectrophotometry. As shown in Table 4, the concentrations of Maxatezo decreased only 3% (from 59.83 mg/mL to 57.93 mg/mL) after the treatments, within the range of loss by filtration. However, the concentrations of aglycosylated Atezolizumab decreased from 60.88 mg/mL to 49.13 mg/mL, which constitute to approximately 20% of loss. Clearly, Maxatezo with GGGS insertion is much more stable than Atezolizumab.

**Table 4.**
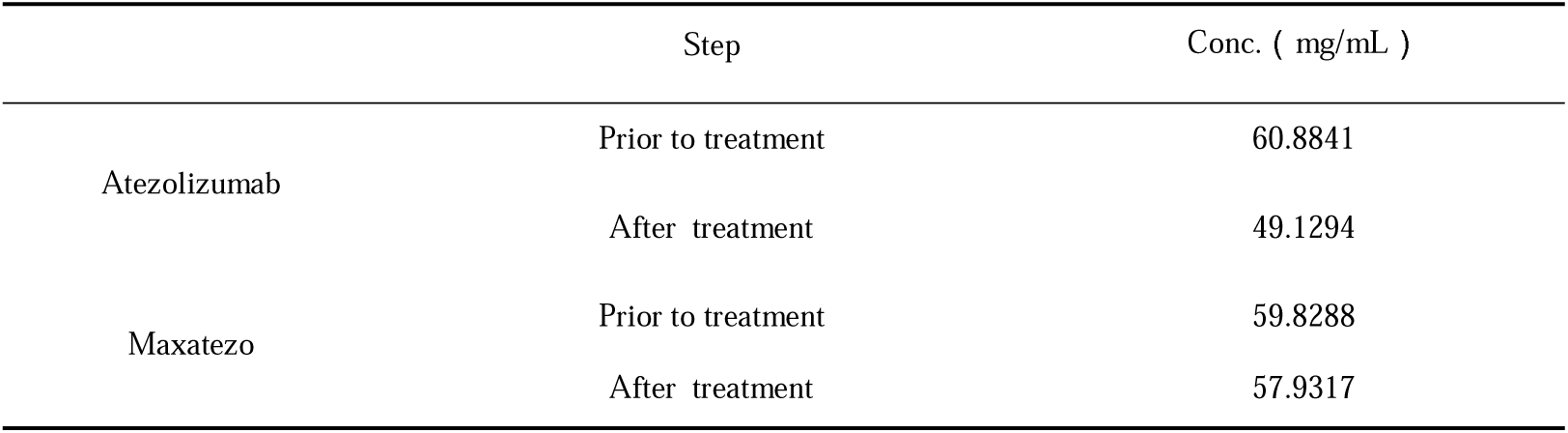
Concentrations of Atezolizumab and Maxatezo after heating

|  | Step | Conc. ( mg/mL ) |
| --- | --- | --- |
| Atezolizumab | Prior to treatment | 60.8841 |
|  | After treatment | 49.1294 |
| Maxatezo | Prior to treatment | 59.8288 |
|  | After treatment | 57.9317 |

**Table 5.** Antitumor Activity of PD-L1 antibodies in the Treatment of MC38 Mouse Colorectal Cancer Model (Tumor Volume)

| Group | Animal No. | Body Weight(g) <sup>a</sup> |  | Tumor Volume(mm<sup>3</sup>)<sup>a</sup> |  | T/C (%) | TGI (%) | *P*<sup>b</sup> | *P*<sup>c</sup> | *P*<sup>d</sup> | Complete tumor regression |
| --- | --- | --- | --- | --- | --- | --- | --- | --- | --- | --- | --- |
|  |  | Before treatment (PG-D0) | After treatment (PG-D17) | Before treatment (PG-D0) | After treatment (PG-D17) |  |  |  |  |  |  |
| Human IgG1 3.3mg/kg | 8 | 28.3±0.6 | 31.7±0.9 | 93±4 | 3912±348 | -- | -- | -- | -- | -- | -- |
| Maxatezo 1mg/kg | 8 | 28.0±0.4 | 29.3±0.4 | 92±4 | 1068±433 | 26% | 74% | 0.000 | 0.018 | 0.361 | 1/8 |
| Maxatezo 3.3mg/kg | 8 | 27.5±0.5 | 29.6±0.8 | 91±4 | 1906±539 | 49% | 51% | 0.000 | 0.000 | -- | 2/8 |
| Maxatezo 10mg/kg | 8 | 28.2±0.5 | 28.4±0.5 | 92±5 | 97±83 | 2% | 98% | 0.000 | -- | 0.000 | 5/8 |
| Atezolizumab 3.3mg/kg | 8 | 28.1±0.7 | 29.1±0.6 | 93±3 | 868±437 | 21% | 79% | 0.000 | 0.155 | 0.051 | 3/8 |
| Atezolizumab 10mg/kg | 8 | 28.4±0.4 | 30.4±1.0 | 93±4 | 1362±784 | 32% | 68% | 0.000 | 0.007 | 0.724 | 4/8 |

Note: a. Mean±SEM; b compared with Human IgG1 3.3mg/kg; c compared with Maxatezo 10mg/kg; d compared with Maxatezo 3.3mg/kg.

*In vivo* immunogenicity assessment was carried out to compare the anti-drug antibody (ADA) titers of Atezolizumab and Maxatezo in mice. As shown in Fig. 5, Atezolizumab induced extremely high titers of ADA in mice, aligning with what was observed in healthy monkeys and human cancer patients. As glycosylation had been restored, the ADA titers of Maxatezo was 70% less than that of the aglycosylated Atezolizumab.

**Figure 5.**
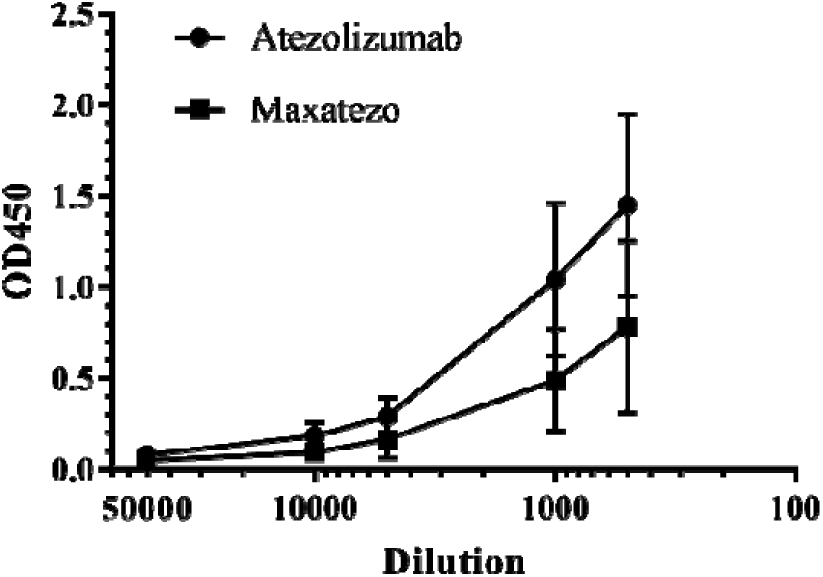
ADA titers evaluation from the mice immunized with either Maxatezo or Atezolizumab. Immunogenicity of antibodies was assessed using *in vivo* mouse model and the ADA titers were measured using Indirect ELISA against antibody drugs. The line with squares represents the Maxatezo, (anti-PD-L1 antibody with GGGS insertion), the line with rhombus represents the Atezolizumab (anti-PD-L1 antibody with N297A). The horizontal coordinate is antibody dilution ratio and the vertical coordinate is the absorbance value at 450nm.

### Anti-tumor Efficacy

In this study, the *in vivo* therapeutic efficacy of the test compounds Maxatezo and Atezolizumab was evaluated in the treatment of MC38 mouse colorectal cancer model in C57BL/6J mice.

As shown in Fig. 6, Maxatezo(1mg/kg, q3d x 6), Maxatezo (3.3mg/kg, q3d x 6), Maxatezo (10mg/kg, q3d x 6), Atezolizumab (3.3mg/kg, q3d x 6) and Atezolizumab (10mg/kg, q3d x 6) treatments produced TGI values of 74%, 51%, 98%, 79% and 68%, respectively. Tumors in treatment groups were all significantly smaller than those in the human IgG1 group (p<0.05). High dose Maxatezo treatment at 10mg/kg resulted in significantly smaller tumors compared with Maxatezo at 1mg/kg and Atezolizumab at 10mg/kg (p<0.05). Complete tumor regression was observed in 1, 2, 5, 3, 4 mice from the Maxatezo groups (1mg/kg, 3.3mg/kg, 10mg/kg) and Atezolizumab groups (3.3mg/kg, 10mg/kg), respectively.

**Fig. 6.**
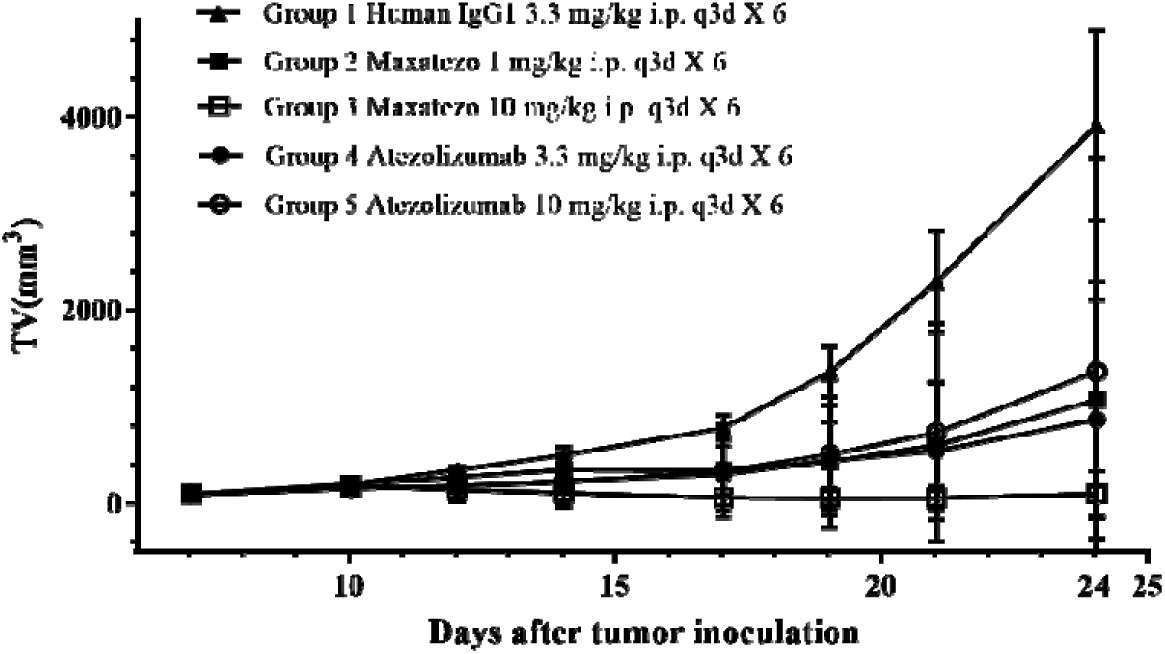
Tumor Growth Curves of Mice in Different Treatment Groups. The horizontal coordinate is the days after tumor inoculation and the vertical coordinate is the tumor volume (mm^3^).

All the mice in this study have been in decent condition without obvious abnormality, dosing holiday or death during the treatment.

In summary, Maxatezo and Atezolizumab produced significant antitumor activity in MC38 mouse colorectal cancer model. Maxatezo (10mg/kg) is significantly better than Atezolizumab to inhibit tumor growth at the same dosing level and frequency of administration. Tumor-bearing mice showed good tolerance to continuous administration of Maxatezo and Atezolizumab in this experiment.

Our data demonstrated that the insertion of GGGS in the hinge regions of human IgG1 could abolish their ADCC activities completely without concering negative impact on antibody affinities, inhibitory activities, expression levels, stabilities or immunogenicity. The efficacy in tumor inhibition of Maxatezo is much better than that of Atezolizumab.

## Discussion

Immunotherapy, using monoclonal antibodies against PD1, PD-L1 or CTLA4, has demonstrated effective to treat various cancers. The therapeutic fields for antibody drugs also expand beyond autoimmune disease, cancers and infectious diseases into chronic diseases such as pain, neurodegenerations, diabetes, and osteoporosis. To reduce the immune-related adverse effects (irAE), switching IgG1 to IgG2/IgG4 or use of aglycosylation of IgG1 has been widely used to remove the Fc function(s) of antibody drugs. However, IgG2 has an extra cystine residue in the upstream of the hinge region. As such, it will form homo- or hetero-dimers via the inter-IgG2 disulfide bond, which will affect its expression level and stability^31^. IgG4 has reduced ADCC but retains ADCP activity. Moreover, it is considered that ADCP activity of Pembrolizumab reduced its tumor killing potency by phagocytosis of NK cells^32^.

A new Fc function removal technology based on structural biology was reported in this paper. A flexible sequence, such as GGGS, was inserted into the hinge region of IgG1 isotype antibody to interrupt the stress signal transfer between the Fv and Fc region. For this reason, the FcγR binding domain will not be exposed when the antibody binds to the target, leading to the loss of ADCC activity.

In this study, the rational design of human IgG1 antibody without ADCC or other Fc functions is a very promising approach to suffice the needs to develop therapeutic antibodies without unwanted antibody’s Fc functions. For those well-known antibody drugs, such as Genentech’s aglycosylated anti-PD-L1 Atezolizumab, we demonstrated that inserting GGGS in the hinge regions of human IgG1 Fc could remove the ADCC activities completely. Since this approach does not alter the Fv domain of the antibody, we did not observe negative impact on either the affinity or inhibitory activity.

The study strategized a physics-based approach to solve a problem of biology. Inspired by previous studies about the structural change of the Fc hinge region after the binding of antigens, we simply inserted a flexible linker to stop the stress transmission from Fv to Fc of the antibody, and to prevent the exposure of binding sites for various FcγRs. Leaving the glycosylation intact and making no additional changes in the remaining parts of the Fc region not only resulted in much higher expression levels than the aglycosylated Atzeolizumab, but stability was also improved. Currently, the re-engineered anti-PD-L1 (Maxatezo) just successfully manufactured and shall proceed into pre-clinical studies shortly.

As of June 2020, there are 10 antibody drugs on the market and more than 50 in clinical trials with purposely reduced ADCC and/or ADCP activities. In separate study with re-engineering anti-PD-1 antibody (Pembrolizumab), anti-CTLA-4 antibody (Ipilimumab) and anti-CD47 antibody (Magrolimab), we also demonstrated that the De-Fc function method could be universal. It may offer a better way to develop safer antibody drugs with less irAE or improve the efficacy of antibody drugs with no ADCC/ADCP. The prospects of this Fc function removal technology are promising.

Furthermore, in the SARS-CoV infection patients, the anti-spike IgG cause severe acute lung injury through the FcγRs, which could skew alveolar macrophages from wound-healing to proinflammatory^33^. We expect that our technology can also help research fellows develop COVID-19 therapeutic antibodies with much less proinflammatory activities.

## Materials and Methods

### Cell lines and reagents

CHO-K1 cells and serum-free medium for transient transfection were obtained from Zhuhai Kairui Biotech, Ltd. The stable cell pool was constructed by Zhenge Biotechnology Co. (Shanghai, China). For stable cell pool construction, the adherent CHO K1 parental cells were obtained from ATCC (American Type Culture Collection, USA). After adapted into suspended and serum-free culture, the CHO K1 cells were cultured in CD CHO medium (Gibco, USA). The viable and total cell counts were determined by ViCell (Beckman, USA). DNA Gel Extraction Kit and Plasmid extraction kit were purchased from Axygen. T4 DNA ligase was from NEB, and Yeast extract and Tryptone were from Oxoid. Sodium chloride and ampicillin were purchased from Amresco. Agar and Agarose were from BioWest and BioSharp. CD CHO was from Gibco, MSX, DMSO Complete Freund’s adjuvant (CFA), incomplete Freund’s adjuvant (IFA), CFSE, PI fluorochrome and TMB substrates were purchased from SIGMA (MO, USA). Female BALB/c mice were obtained from Vital River Co. (Beijing, China). Goat anti-mouse IgG Fc and Goat anti-human IgG secondary antibodies are from Jackson Immune Lab (MA, USA). Recombinant PD-1, PD-L1 and CTLA-4 were purchased from Sino Biological (Beijing, China). The CHO-PD-L1-CD3L cell and Jurkat-PD-1-NFAT cell line were provided by National Institutes for Food and Drug Control (short for NIFDC) (Beijing, China). FcR-TANK cells and target protein (such as PD-1, PD-L1 or CTLA-4) overexpressing Raji cells were provided by ImmuneOnco Biopharma Co., Ltd (Shanghai, China).

### Antibody production

The antibodies were expressed either by transient transfection or stable cell pool, and CHO-K1 cells were used for both routes. All the antibodies were purified with Protein A Sepharose.

For the transient transfection, the heavy chain of antibody was constructed into pEE12.4 plasmid, while the light chain of antibodies was constructed into pEE6.4 plasmid. For antibody transient expression, CHO-K1 cells were co-transfected with the plasmids containing either heavy chain or light chain at the ratio of 1:1. And the transfected cells were cultured at 37 □, 5% CO_2_ for 6 days. Then the supernatants were collected, and the antibodies were purified with Protein A Sepharose.

To establish the stable cell pool, after the transfection, CHO host cells were subjected to selection pressure to obtain stable mini cell pools, and the antibody expression level by fed batch in 2L bioreactor were evaluated. To obtain monoclonal stable cell line, sub-cloning by limiting dilution was carried out from the selected stable mini pools. Top 8-10 clones were selected for further development.

### Indirect ELISA

The binding affinity was determined by Indirect ELISA against the antigen. Each well of the 96-well high binding EIA plates was coated with 1 µg/mL of antigen, such as recombinant PD-L1, at 4□ overnight in PBS. After two washes with PBS and blocking with 5% skim-milk in PBS for 1 hour at room temperature, wells were incubated with purified antibody in 5% skim-milk-PBS for another one hour at room temperature. After two washes with PBS, wells were then incubated with HRP-conjugated goat anti-human IgG Fc-specific secondary antibodies (Jackson Lab) in 5% skim-milk-PBS for 1 hour at room temperature. After five washes with PBS plus 0.1% Tween20 (PBST), HRP substrate 3, 3’, 5, 5’-tetramethylbenzidine (TMB) solution was added. The reaction was stopped with stop solution (0.1M H_2_SO_4_) after 30 minutes and absorbance was measured at 450nm with a microplate reader.

### Bioassay of anti-PD-L1 antibodies

As reported previously^34^, two cell lines were used for this assay: the CHO-PD-L1-CD3L cell line that stably over-expressing PD-L1 and a membrane-anchored anti-CD3 single chain antibody fragment (scFv), and the Jurkat-PD-1-NFAT cell line that stably over-expressing PD-1 and the luciferase gene under the control of the NFAT responses elements from the IL-2 promoter.

Briefly, Jurkat-PD-1-NFAT cell line that stably expresses PD-1 with the reporter luciferase gene were added to the wells cultured with CHO-PD-L1-CD3L cells that stably expresses PD-L1 and a membrane-anchored anti-CD3 single chain antibody fragment (scFv). The binding of PD-1 on the surface of Jurkat-PD-1-NFAT cells to the PD-L1 on the CHO-PD-L1-CD3L cells can block the downstream signal transduction of CD3 activated by the binding of anti-CD3 scFv, thus inhibiting the expression of luciferase in Jurkat-PD-1-NFAT cells. When PD-1 antibody or PD-L1 antibody is added, it will reverse the inhibition of luciferase expression.

CHO-PD-L1-CD3L cells were seeded at 50,000 cells per well in 96 well plate, incubated at 37□, with 5% CO2 for 12-14 hours. 100,000 of Jurkat-PD-1-NFAT cells were added to each well in the presence or absence of testing antibody and incubated at 37□ with 5% CO2 for another 6 hours. Cells were lysed and 100 μL of luciferase substrate (Promega Bio-GloTM Luciferase Assay) was added into each well, and the plate was measured using SpectraMax M5 to calculate the relative luciferase unit.

### *In vitro* ADCC detection assay

Using CFSE to dye Raji cells overexpressing target protein (such as PD-L1, PD-1 or CTLA-4), then mixed with FcR-TANK cell in the presence or absence of testing antibody, incubated at 37□, with 5% CO_2_ for 4 hours. At the end of incubation, add 5μg/mL PI was added to stain the dying Raji cells, and the cell populations were analyzed using flow-cytometer. For calculation of ADCC activity: ADCC% = [(Sample % PI Positive cell – Control % PI Positive cell) / (100 – Control % PI Positive cell)] × 100.

### HPLC

For HPLC-SEC analysis of antibodies, the sample was diluted to 2.0 mg/mL with the mobile phase (50 mM NaH_2_PO4, 300 mM NaCl, pH 7.0), then loaded 50μL to the TSK gel G3000SWXL (5μm, 7.8×300 mm) column which has been equilibrated by the same mobile phase. Then run the procedure with 0.5 ml / min flow rate for 90 min. The absorbance value at 280 nm was monitored.

### Glycan assay

Antibodies were first digested with PNGase F (New England Biolabs) according to the manufacturer’s instructions, and the free glycan(s) were separated using LudgerClean™ EB10 kit according to the manufacturer’s instructions. The glycan samples were labeled with 2-AB (2-aminobenzamide) using LudgerTag™ 2-AB (2-aminobenzamide) Glycan Labeling Kit and separated with LudgerClean™ S cartridges. The 2-AB labeled glycan samples were analyzed by HPLC with fluorescence detection.

### *In vivo* immunogenicity assay

As reported previously^35^, Balb/c mouse were used to evaluate the ADA titers of different antibodies. 4∼6 weeks old female Balb/c mice were first immunized with antibodies in Complete Freund’s Adjuvant and boosted with antibodies in Incomplete Freund’s Adjuvant. Two to four weeks after the first immunization, tail bleeds from the immunized mice were tested for titers by indirect Enzyme-linked Immunoassay (ELISA) against antibody drugs in the presence of 1% human sera.

### *In vivo* Efficacy Study of Maxatezo and Atezolizumab in MC38 Mouse Colorectal Cancer Model

The MC38 cell line purchased from Biovector NTCC Inc.. The tumor cell is maintained *in vitro* in DMEM medium supplemented with 10% heat inactivated fetal calf serum, 100U/ml penicillin, 100 μg/ml streptomycin at 37°C in an atmosphere of 5% CO_2_ in air. The cells growing in an exponential phase are harvested and counted for tumor inoculation.

Tumor cells were suspended in PBS to 1×10^7^/ml after two washes and they were subcutaneously inoculated at the right flank of C57BL/6J mice with 100µl/mouse. When the average tumor volume reached about 93mm3, the mice were randomized based on their tumor volumes and test articles were administered to the mice according to the predetermined regimen as shown in Table S2.

After inoculation, the animals were checked daily for morbidity and mortality. At the time of routine monitoring, the animals were checked for any effects of tumor growth and treatments on normal behavior such as mobility, food and water consumption, body weight gain/loss, eye/hair matting and any other abnormal effect. Animal death and observable clinical signs will be recorded.

The tumor volume was calculated according to formula: tumor volume = 0.5 × long diameter × short diameter^2^, and the tumors were measured twice a week. The T/C values were calculated from the tumor volume, where T is the average relative tumor volume (RTV) of each test subject treated group, and C is the average RTV of the control group. RTV is the ratio of tumor volume after administration to pre-dose. Tumor growth inhibition (TGI%) was calculated as (1-T/C) × 100%. The treatments with tumor growth inhibition ≥ 60% and statistically significant difference in tumor volume are considered effective.

Tumors were collected, photographed, weighed at study termination and tumor weight inhibition (%) was calculated as (1-average TW of each test subject treated group/average TW of the control group) × 100%.

