## Supplemental Figure for "Next generation of anti-PD-L1 Atezolizumab with better anti-tumor efficacy *in vivo*"

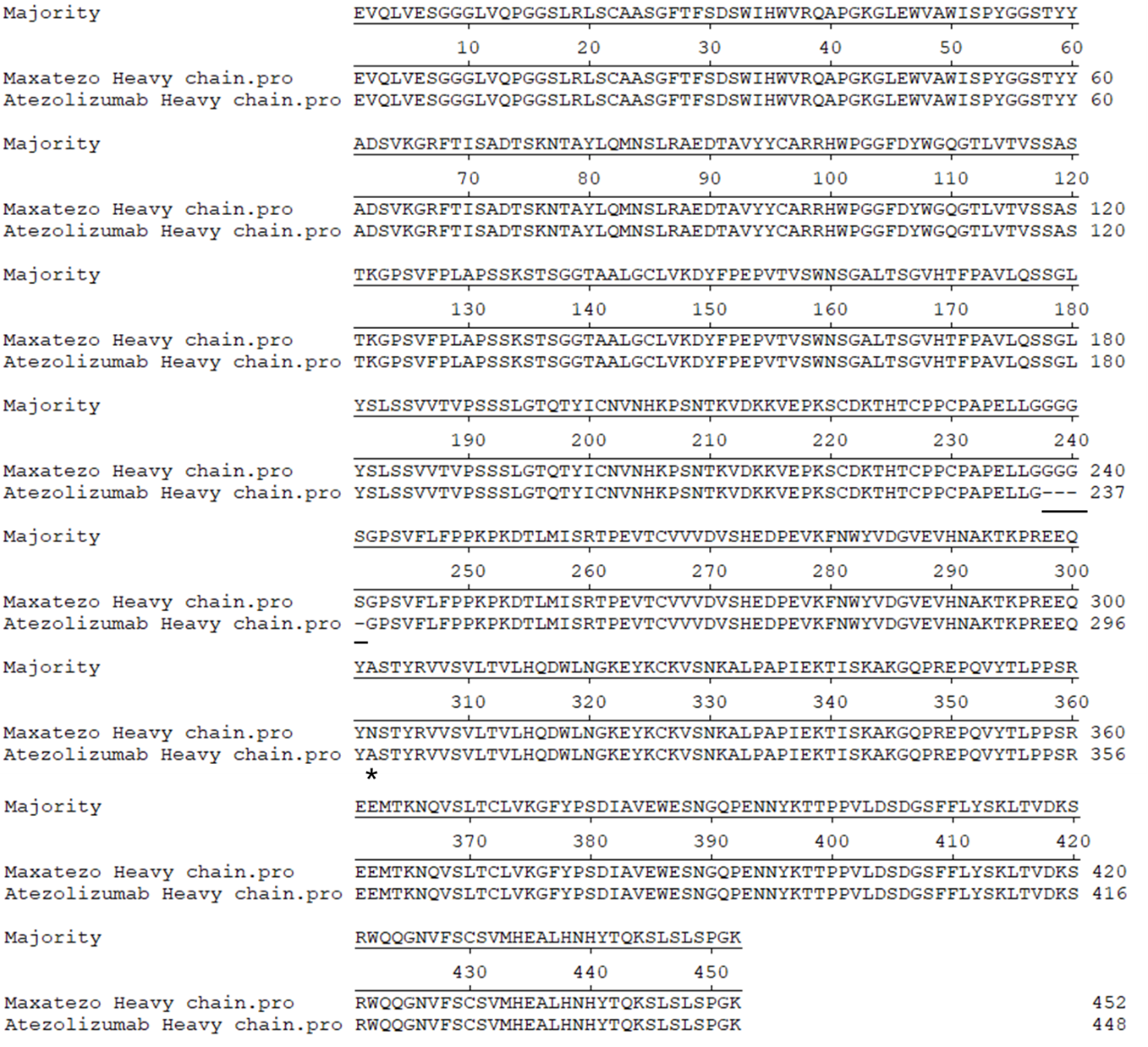

**Figure S1. Sequence alignment of atezolizumab and maxatezo**

(a). Sequence alignment of Atezolizumab and Maxatezo. The protein sequences of Atezolizumab and Maxatezo were aligned by clustal W method. The insertion sequence and site were marked by underline. The back mutation of A297N was indicated by *.

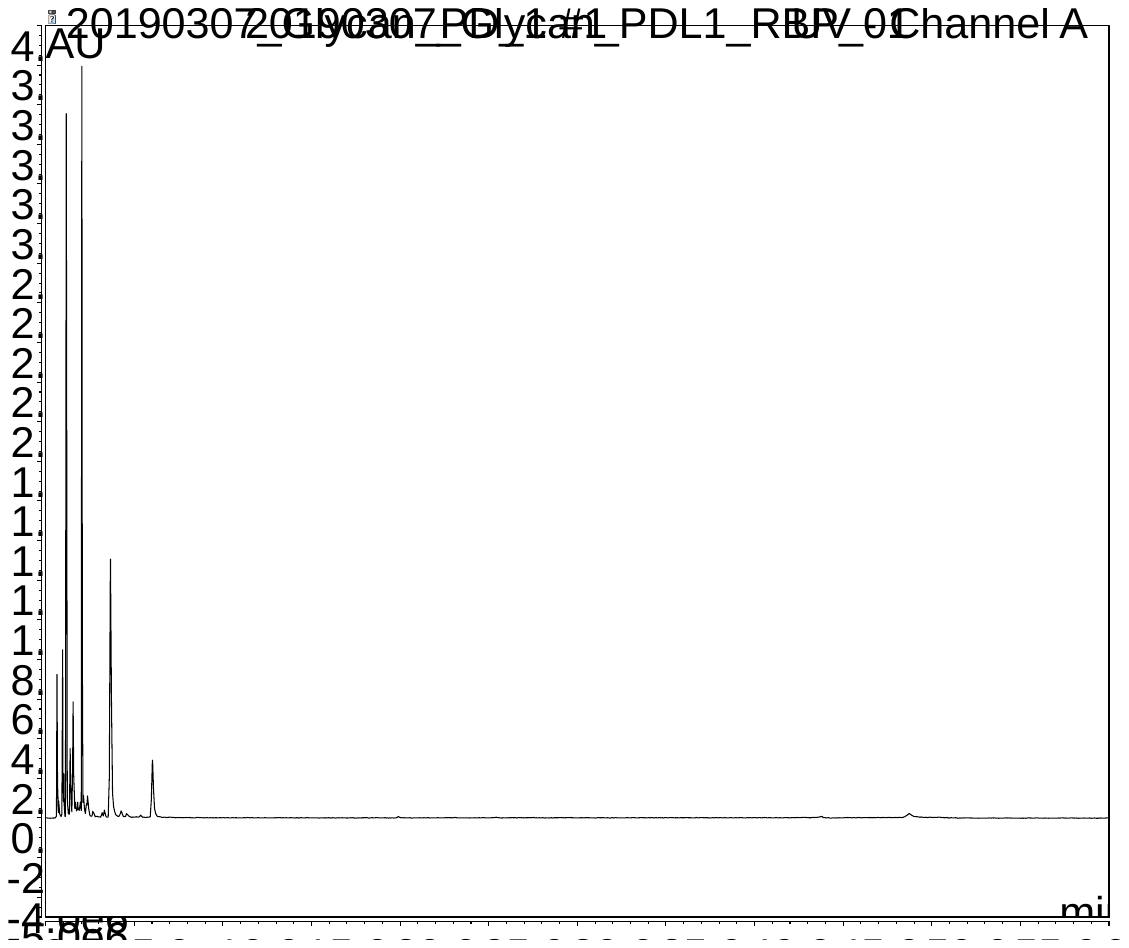

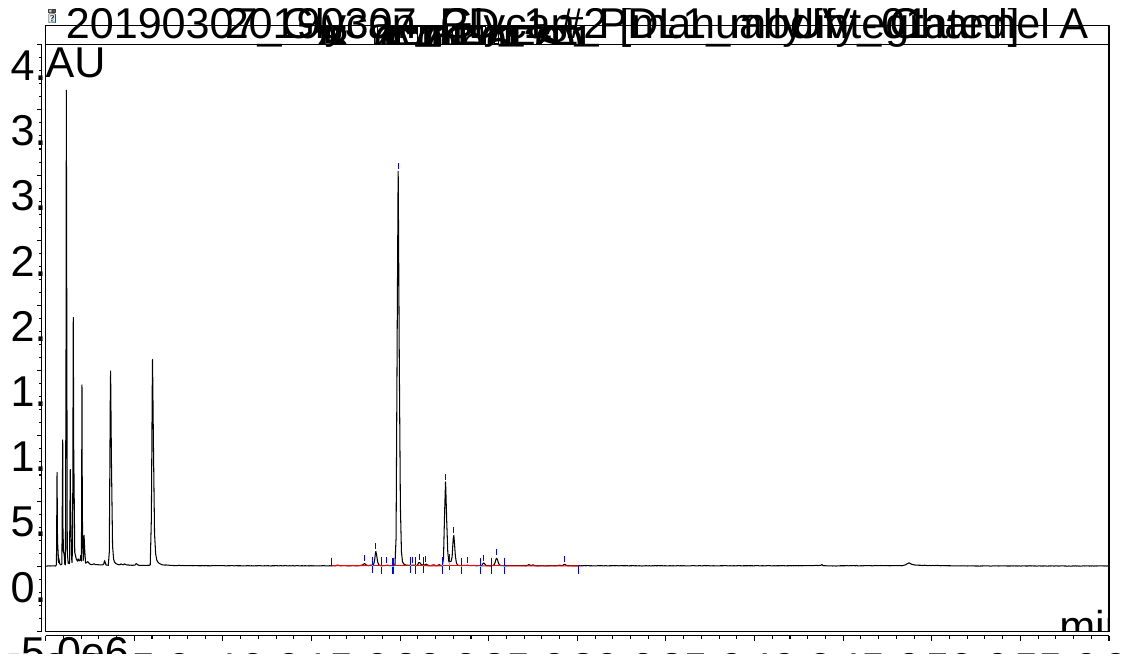

**Figure S2. Glycan analysis of anti-PD-L1 mAbs.**

. (A) The HILIC of Atezolizumab; (B) The HILIC of Maxatezo. The horizontal coordinate is time and the vertical coordinate is absorbance value at 280nm.

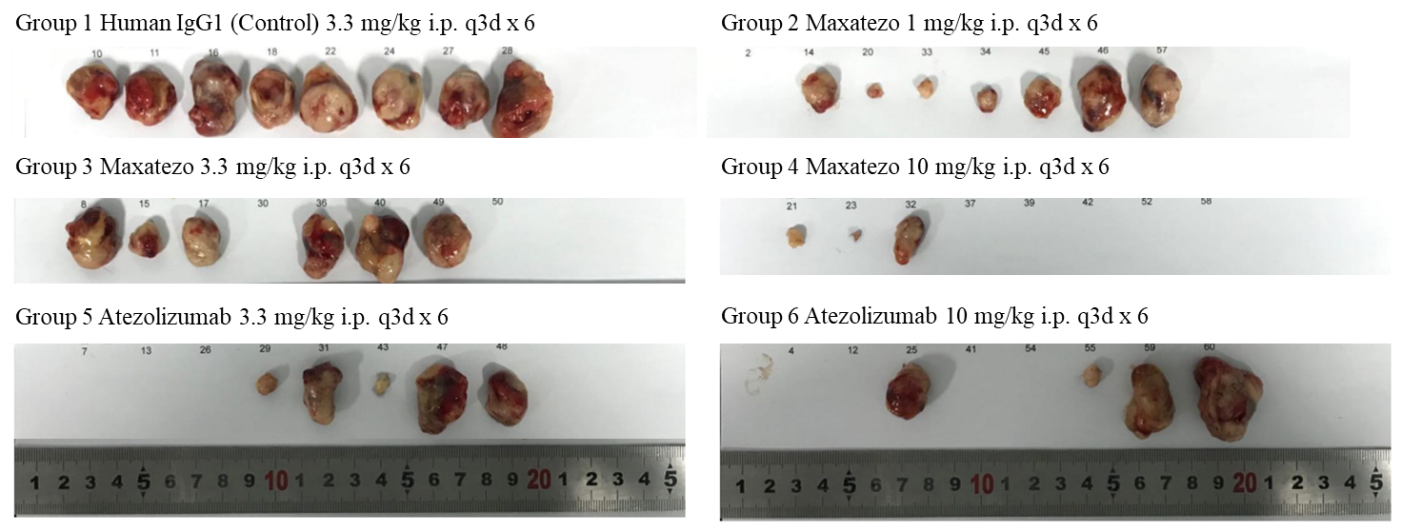

**Fig. S3 Photograph of tumors at the termination point**

The *in vivo* therapeutic efficacy of the Maxatezo and Atezolizumab was evaluated in the treatment of MC38 mouse colorectal cancer model in C57BL/6J mice. The mice were randomized based on their tumor volumes and test articles were administered with differernt dose. Tumors were collected, photographed, weighed at study termination.
